# Neurodesk: An accessible, flexible, and portable data analysis environment for reproducible neuroimaging

**DOI:** 10.1101/2022.12.23.521691

**Authors:** Angela I. Renton, Thuy T. Dao, Tom Johnstone, Oren Civier, Ryan P. Sullivan, David J. White, Paris Lyons, Benjamin M. Slade, David F. Abbott, Toluwani J. Amos, Saskia Bollmann, Andy Botting, Megan E. J. Campbell, Jeryn Chang, Thomas G. Close, Korbinian Eckstein, Gary F. Egan, Stefanie Evas, Guillaume Flandin, Kelly G. Garner, Marta I. Garrido, Satrajit S. Ghosh, Martin Grignard, Anthony J. Hannan, Renzo Huber, Jakub R. Kaczmarzyk, Lars Kasper, Levin Kuhlmann, Kexin Lou, Yorguin-Jose Mantilla-Ramos, Jason B. Mattingley, Jo Morris, Akshaiy Narayanan, Franco Pestilli, Aina Puce, Fernanda L. Ribeiro, Nigel C. Rogasch, Chris Rorden, Mark Schira, Thomas B. Shaw, Paul F. Sowman, Gershon Spitz, Ashley Stewart, Xincheng Ye, Judy D. Zhu, Matthew E. Hughes, Aswin Narayanan, Steffen Bollmann

**Affiliations:** The University of Queensland, Queensland Brain Institute, St Lucia 4072, Australia; The University of Queensland, School of Information Technology and Electrical Engineering, St Lucia 4072, Australia; Medical Scientist Training Program, Stony Brook University, Stony Brook, NY, United States of America; Discipline of Psychiatry, Adelaide Medical School, University of Adelaide, Australia; The University of Queensland, Centre for Advanced Imaging, St Lucia 4072, Australia; The University of Queensland, School of Psychology, St Lucia 4072, Australia; Centre for Mental Health & Brain Sciences, Swinburne University of Technology, Hawthorn 3122, Australia; School of Psychology, University of Adelaide, Adelaide, 5000, Australia; Human Health, Health & Biosecurity, CSIRO, Adelaide, 5000, Australia; Macquarie University, School of Psychological Sciences, North Ryde 2112, Australia; Department of Neurology, Royal Brisbane and Women's Hospital, Brisbane, Australia; Hopwood Centre for Neurobiology, Lifelong Health Theme, South Australian Health and Medical Research Institute (SAHMRI), Adelaide, SA, Australia; The Turner Institute for Brain and Mental Health, School of Psychological Sciences, Monash University, Victoria, Australia; Simons Center for Quantitative Biology, Cold Spring Harbor Laboratory, Cold Spring Harbor, NY, United States of America; The Florey Institute of Neuroscience and Mental Health, The University of Melbourne, Victoria, Australia; The University of Sydney, School of Biomedical Engineering, Sydney, Australia; Monash Biomedical Imaging, Monash University, Victoria, Australia; School of Psychological Sciences, University of Newcastle, Australia; Hunter Medical Research Institute Imaging Centre, Newcastle, Australia; Functional Magnetic Resonance Imaging Core Facility (FMRIF), National Institute of Mental Health (NIMH), USA; Techna Institute, University Health Network, Toronto, Canada; Melbourne School of Psychological Sciences, The University of Melbourne; Graeme Clark Institute for Biomedical Engineering, The University of Melbourne; Department of Neuroscience, Central Clinical School, Faculty of Medicine, Nursing and Health Sciences, Monash University, Australia; The University of Queensland, School of Biomedical Sciences, St Lucia 4072, Australia; Department of Psychology, Center for Perceptual Systems, Center for Theoretical and Computational Neuroscience, Center on Aging and Population Sciences, Center for Learning and Memory, The University of Texas at Austin, 108 E Dean Keeton St, Austin, TX 78712, USA; Department of Data Science and AI, Faculty of Information Technology, Monash University, Clayton VIC 3800, Australia; Department of Psychological and Brain Sciences, Indiana University, Bloomington, IN 47405, USA; Monash-Epworth Rehabilitation Research Centre, Turner Institute for Brain and Mental Health, School of Psychological Sciences, Monash University, Clayton, 3168, Australia; McGovern Institute for Brain Research, Massachusetts Institute of Technology, Cambridge, MA 02139, USA; Department of Otolaryngology - Head and Neck Surgery, Harvard Medical School, Boston, MA, USA; GIGA CRC In-Vivo Imaging, University of Liège, Liège, Belgium; Wellcome Centre for Human Neuroimaging, University College London, London, UK; School of Life Science and Technology, University of Electronic Science and Technology, China; Grupo Neuropsicología y Conducta (GRUNECO), Facultad de Medicina, Universidad de Antioquia, Colombia; Australian Research Data Commons (ARDC), Australia; McCausland Center for Brain Imaging, Department of Psychology, University of South Carolina, Columbia SC, 29208, USA; The University of Auckland, Auckland, 1010, New Zealand; School of Psychology, University of Wollongong, Wollongong, 2522, Australia

## Abstract

Neuroimaging data analysis often requires purpose-built software, which can be challenging to install and may produce different results across computing environments. Beyond being a roadblock to neuroscientists, these issues of accessibility and portability can hamper the reproducibility of neuroimaging data analysis pipelines. Here, we introduce the Neurodesk platform, which harnesses software containers to support a comprehensive and growing suite of neuroimaging software (https://www.neurodesk.org/). Neurodesk includes a browser-accessible virtual desktop environment and a command line interface, mediating access to containerized neuroimaging software libraries on various computing platforms, including personal and high-performance computers, cloud computing and Jupyter Notebooks. This community-oriented, open-source platform enables a paradigm shift for neuroimaging data analysis, allowing for accessible, flexible, fully reproducible, and portable data analysis pipelines.

## Introduction

Neuroimaging data analysis is a challenging enterprise. Aside from the neuroscientific principles motivating the choice of analysis, building an analysis pipeline requires advanced domain knowledge well beyond the researcher’s topic area; for example, signal and image processing, computer science, software engineering, statistics, machine learning, and applied physics. Researchers faced with this daunting task rely on multiple specialized software packages used in custom pipelines to suit a specific aim. Researchers with limited resources and software engineering teams often develop these packages, resulting in little dedicated technical support. The required software packages are often difficult to install and are inconsistently supported across computing environments^2–4^. They often have conflicting dependencies tied to the specific operating system versions. Consequently, researchers often spend considerable time installing and compiling bespoke neuroimaging software, which can undermine scientific productivity and reproducibility. To address these issues, we developedan open-source and community-oriented solution to enable neuroscientists to develop neuroimaging analysis workflows in line with four guiding principles: *Accessibility*, *Portability*, *Reproducibility*, and *Flexibility*.

Ideally, the software and code used in any scientific analysis workflow should be easily *accessible* so that users can deploy the workflow without a substantial investment of time or effort^5^. It should be *portable* so that analysis workflows can be tractably shifted between operating system versions and computing environments and deliver identical results. Many researchers prototype analysis pipelines using their local computers and later switch to workstations and high-performance computing (HPC) clusters for processing datasets at scale. Accessible and portable workflows allow for an optimized allocation of computing resources while supporting shared development workloads amongst collaborators^6^. Unfortunately, many neuroimaging data analysis workflows are currently neither readily accessible nor portable^7–9^ because they rely on specialized tools purpose-built by a small number of developers^2^.

Beyond the productivity costs, the inaccessibility and instability of many neuroimaging tools pose a wider threat to *reproducibility*^10–17^ with reproducibility defined as “running the same software on the same input data and obtaining the same result”^16,18,19^. The transparency and openness promotion (TOP) guidelines, which have over 5,000 journals and organizations as signatories, state that all reported results should be independently reproduced before publication^20^. However, this is impractical and too time-consuming to implement at review^8^. Where analysis pipelines are ported, subtle differences in the implementation of specific processing steps and software versions across computing environments can systematically affect results^21–24^. Thus, it is often impossible to reproduce a prior study’s results, even given the original data and analysis protocol^14,21^. Controlling the specific software version of a tool and its dependencies is key to reproducibility^25^.

Unfortunately, many existing solutions lack the required *flexibility* for research applications of neuroimaging data analysis^26^. For example, single-install pre-programmed analysis pipelines are a popular solution amongst clinicians, but researchers typically custom-tailor analysis pipelines toward specific research questions^27–29^. The issues of inaccessibility in neuroimaging software have been recognized by the NeuroDebian^2^ and NeuroFedora^30^ projects, which provide a wide range of neuroimaging tools packaged for Linux operating systems. However, most neuroscientists do not use Linux on their personal computers and thus cannot access these packages^3^. Researchers often use dual-boot computers or virtual machines to address this barrier. Still, these solutions are resource intensive and force researchers to develop inflexible workflows due to the practical limitations inherent in installing new tools. While compiled packages make installations easier, applications still need to be installed on the host computer and suffer the usual problems of conflicts between different software packages, software versions, or the required libraries (software “dependencies”). Many researchers are also limited in flexibility by institutional restrictions imposed on the installation of new software.

Applications with specific or conflicting dependencies are not unique to neuroscience. This universal issue has led to the development of software containers: lightweight, portable solutions for running and sharing individual applications. Software containers package specific applications along with their dependencies. Container engines such as Docker and Apptainer/Singularity allow applications to run on various computing environments while eliminating concerns about conflicting or missing dependencies^31,32^. These benefits make software containers suited to tackle issues relating to developing the scientific analysis workflows described above^33^. However, despite the benefits of containerization, only a small number of integrated neuroscience-specific or adaptable workflow systems support containerized distributed computing^6,34–36^. Although projects such as OpenNeuro^37^, Brainlife^38^, Flywheel^39^, Boutiques^40,41^, XNAT^42^ and Qmenta^43^ have improved the accessibility and reproducibility of neuroimaging analyses, these projects still lack portability. At this stage, no solution exists that universally addresses the issues of *Accessibility*, *Flexibility*, *Portability,* and *Reproducibility*. Our objective is to change this with the development of Neurodesk: a community-oriented open-source platform that harnesses software containers to create an accessible and portable data analysis environment that allows users to flexibly and reproducibly access a comprehensive range of neuroimaging tools.

## Results

### Overview of the Neurodesk Platform

Here, we present Neurodesk, a platform facilitating *Accessibility, Portability, Reproducibility,* and *Flexibility* for Neuroimaging data analysis (**Figure 1**). In developing Neurodesk, we ensured that workflows developed on the Neurodesk platform remained consistent with these four guiding principles across updates to users’ local computing environments. In this section, we introduce the available tools in the Neurodesk platform, discuss how this addresses the issues raised above and report the results of an empirical evaluation of reproducibility in Neurodesk. For further details of the rationale behind the approaches adopted to achieve these results, please see the online methods.

**Figure 1.**
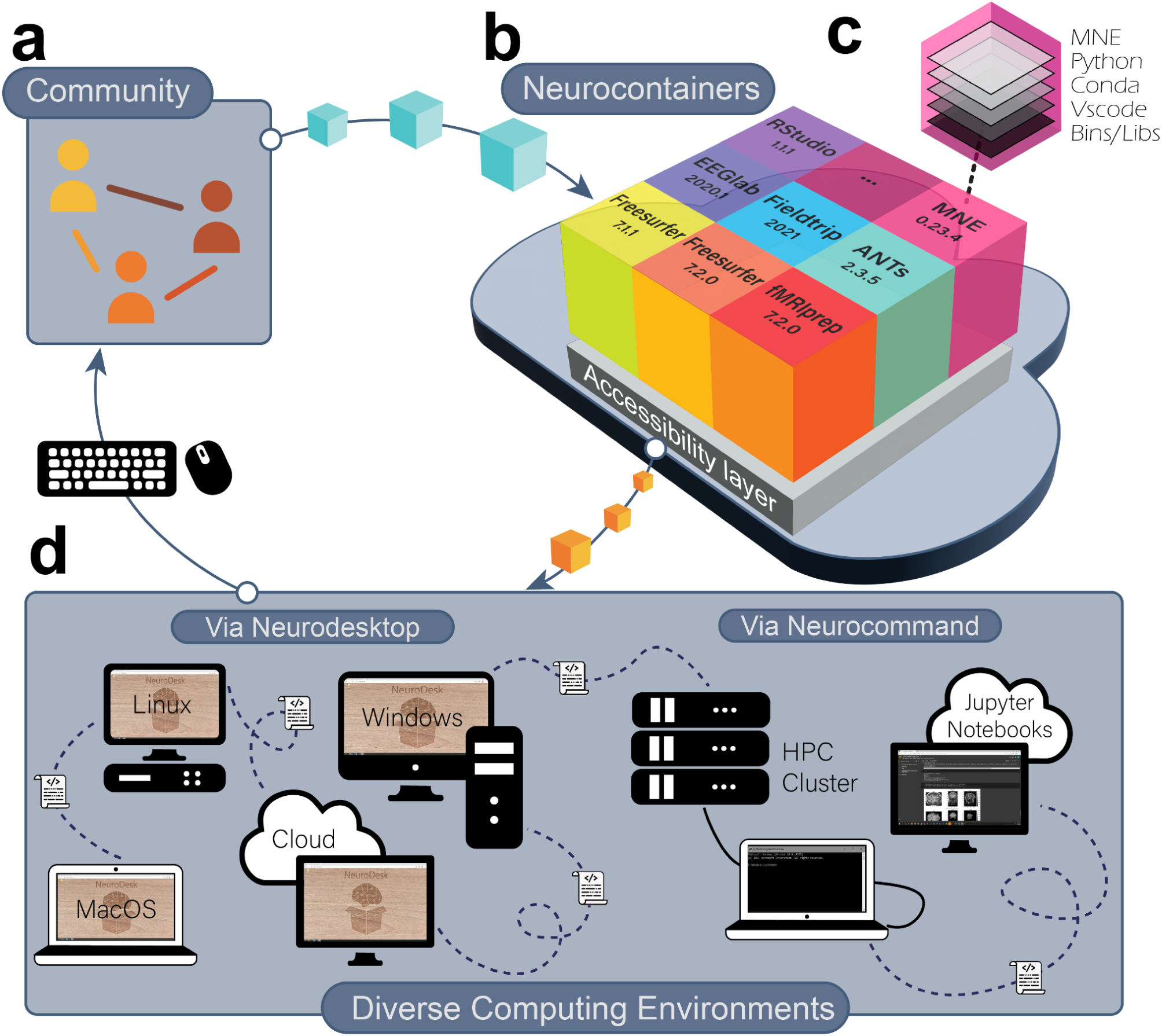
The Neurodesk platform. (**a**) The Neurodesk platform is built by and for the scientific community, enabling anyone to contribute recipes for new software containers to the repository. (**b**) Recipes contributed by the community are automatically used to build software containers and stored in the Neurocontainers repository. (**c**) Each software container packages a tool together with all the required runtime dependencies. The packaged software can therefore run identically in any supported computing environment. (**d**) Neurodesk provides two layers of accessibility: 1. Neurodesktop is a browser-accessible virtual desktop environment, allowing users to interact with the containerized software. 2. Neurocommand is a command-line interface that allows users to run the same software containers programmatically. These interfaces allow users to reproduce the same analysis pipelines across various computing environments.

At the core of Neurodesk are Neurocontainers, a collection of software containers that package a comprehensive and growing array of versioned neuroimaging tools (**Figure 1*b***). The community contributes recipes based on the open-source project Neurodocker^44^, a continuous integration system builds the containers and uploads them to a container registry (**Figure 1*a***). Each ‘Neurocontainer’ includes the packaged tool and all dependencies required to execute that tool, allowing it to run on various computing systems (**Figure 1*c***). Because the containers isolate dependencies, different Neurocontainers can provide different versions of the same tool without conflicts. This mechanism allows researchers to seamlessly transition between different software versions across projects or within a single analysis pipeline. A newly developed accessibility layer enables researchers to use software directly through the cloud or download containers for offline use without the need to install software on a local system (**Figure 1*b***).

There are two options for interfacing with Neurocontainers: The first is Neurodesktop, a remote desktop and browser-accessible virtual desktop environment that can launch any of the containerized tools from the application menu (**Figure 1*d***). Analyzing neuroimaging data through Neurodesktop has the look and feel of working on one’s local computer. For more advanced users and HPC environments, Neurocommand enables interfacing with Neurocontainers through the command line (**Figure 1*d***). These interfaces can be deployed across almost any computing hardware and modern operating system, meaning that analysis pipelines developed using the Neurodesk platform are reproducible and can range from local computers to cloud and HPC environments. Neurocontainers even work inside Jupyter Notebooks, so researchers developing analysis pipelines using Neurodesk can share the reproducible code and results alongside published manuscripts (**Figure 1*d***).

### How to use Neurodesk: Accessibility, Flexibility & Portability

A core aim behind Neurodesk is to provide a platform that makes building and running reproducible analysis pipelines accessible to all researchers. The platform website (https://Neurodesk.org/) is user-friendly and open to community contributions via pull requests. The website contains automatically updating information about the software included through continuous integration. As such, there is always up-to-date documentation, lists of currently available applications, and release history. The website also hosts clear instructions and guidance for accessing and interacting with Neurodesk from various computing environments and tutorials on using various software packages.

Besides ensuring that users have access to thorough and up-to-date documentation, additional steps ensure that Neurodesk makes reproducible neuroimaging data analysis *accessible*. Neurodesk works in almost any computing environment and brings the same dependencies to all supported platforms. This portability extends to the Neurodesktop graphical user interface (GUI), which provides the same desktop environment across all supported computing environments. Containerized analyses look, feel, and run the same way across different computing environments. Thus, researchers reading or reviewing manuscripts with open-source data and code can use Neurodesk to replicate the *exact* pipeline using the reported tool versions without requiring to install additional software.

For a data analysis environment to be *portable*, such that it can easily shift between computing environments, it also needs to be lightweight with a small storage footprint. To this end, our accessibility layer harnesses the CernVM File System (CVMFS)^45^. The CVMFS layer allows accessing the software from a remote host without installation, so only those parts of a container actively used are sent over the network and cached on the user’s local computer. Users can access terabytes of software without downloading or storing it locally. The Neurodesk platform has several CVMFS nodes worldwide, providing low latency and direct access to Neurocontainers. Thus, to use Neurodesk, users only install the required container engine and access the Neurocontainer of their choice. For Neurodesktop, which facilitates access to all tools in the Neurocontainers repository, the download is only ∼1GB.

Anticipating that installing a third-party container engine software may be a barrier to entry for some researchers, there is an entirely cloud-based solution; ‘Neurodesk Play’ (http://play.neurodesk.org). Neurodesk Play is accessible globally, allowing anyone to use a cloud-based graphical desktop environment for neuroimaging data analysis and teaching. Neurodesktop can also run on institutional or cloud computing resources enabling access to large amounts of computing resources or datasets. For example, Neurodesk is freely available as a national service on the Nectar Research Cloud Virtual Desktop Service provided by the Australian Research Data Commons (ARDC).

### Long-Term Sustainability of the Neurodesk Platform

Neurodesk has a wide selection of tools available spanning many domains of neuroimaging data analysis. **Table 1** shows the tools available at the time of publication, though this list is growing rapidly. Users can find a full and up-to-date list at https://Neurodesk.org/applications/. Neurodesk employs a two-pronged approach to staying up-to-date with new neuroimaging tools and new versions of already included software: a.) The Neurodesk maintainers add tools as they become aware of new developments or community members request the addition of new packages. The Neurodesk GitHub repository (https://github.com/NeuroDesk) has an active discussion forum where developers respond to requests for new software containers. b.) In addition to this developer-centric route to new software containers, we actively encourage contributions from the research community. A core aim for developing the Neurodesk platform was to build a community-driven project that is not contingent on a specific team of developers. As such, we provide a template and detailed instructions for creating build scripts for new software containers.

**Table 1.**
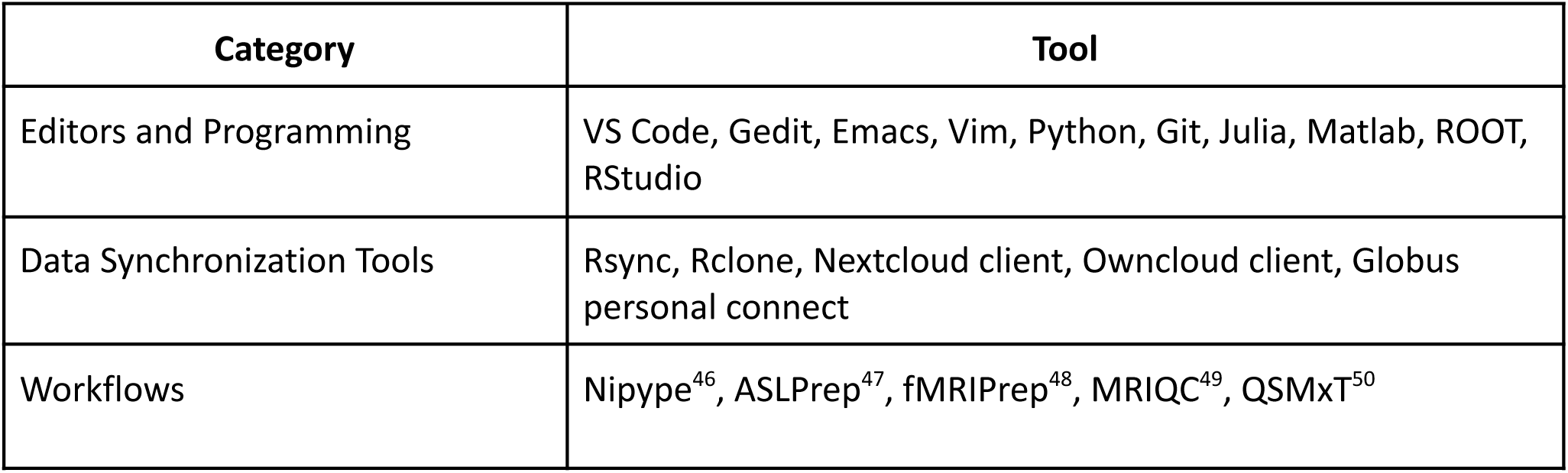
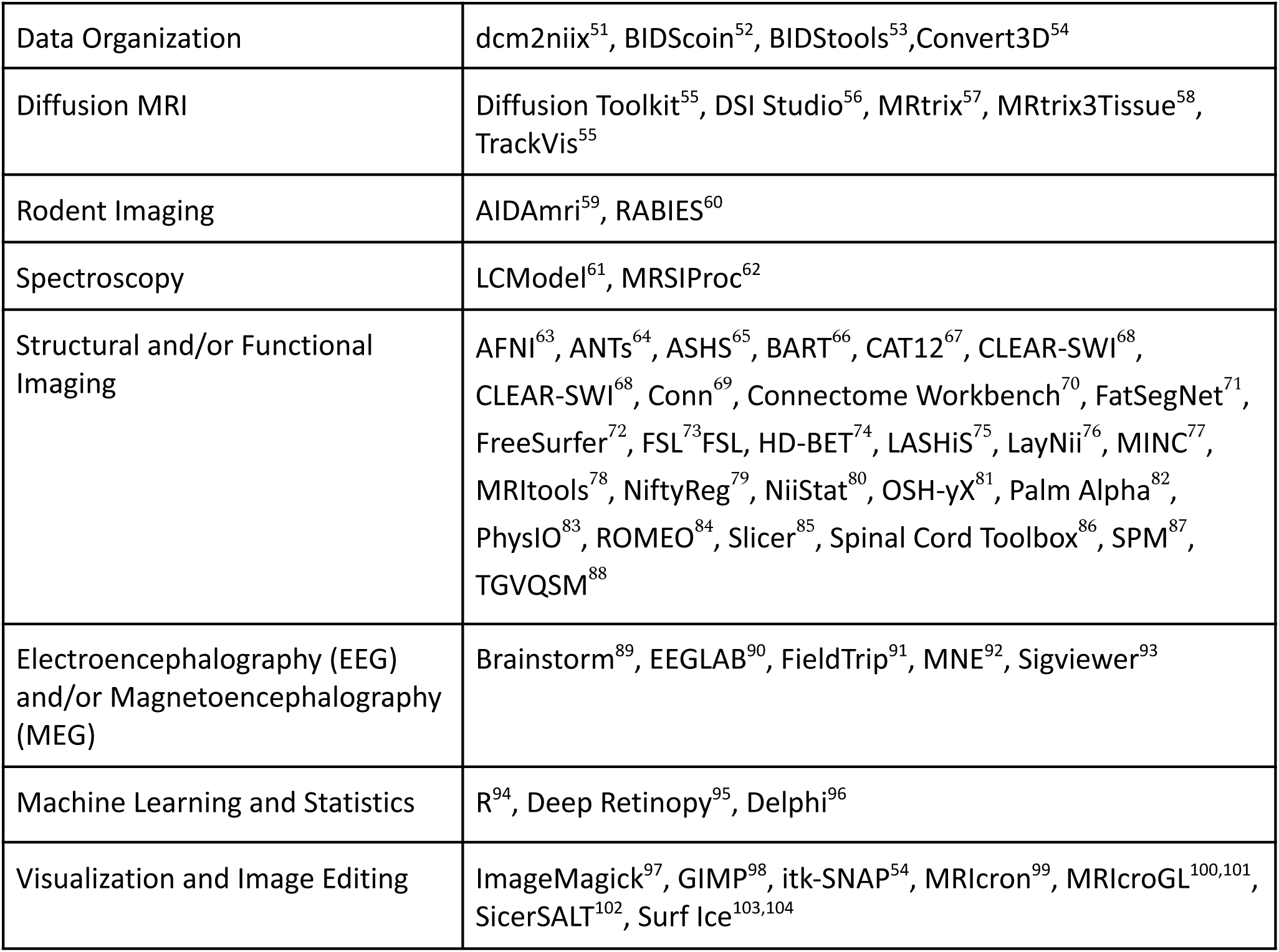
Tools currently available in Neurodesk (retrieved from https://Neurodesk.org/applications/). Note that each tool has been listed under only one category, though some may span multiple categories.

### Reproducibility in Neurodesk

Scientific progress fundamentally depends on the peer review process - scientists must be able to critically assess reported findings and conclusions based on a clear and thorough methodological description^18^. Well-documented experimental code is the most thorough description of any analysis pipeline. However, differences in computing environments and dependencies mean that access to this source code does not guarantee the capability to run the code nor the same result^19,105^. Reproducibility has therefore come to represent a minimum standard by which to judge scientific claims^16,18,19^. Unfortunately, scientific reproducibility is often not attainable due to differences in the outcomes of neuroimaging pipelines across different computing environments, as previously documented^21,106,107^. Glatard et al. (2015) demonstrated this effect for several MRI analysis pipelines, showing that differences in the implementation of floating-point arithmetic across operating systems accumulated throughout long analysis pipelines and led to meaningful differences in the results^21^. Neurodesk solves this issue using containerized software, which guarantees the same runtime dependencies across computing environments. To evaluate this claim, we replicated Glatard et al.’s analyses using Neurodesk vs. locally installed software across different operating systems. We discuss in the online supplements why differences between library versions affect the results.

#### Methodological approach

The widely used FMRIB Software Library (FSL) 6.0.5.1^73^ was installed both locally and within Neurodesk on two separate computers (System A, System B) which were running different Linux distributions. This resulted in four unique computing environments (see **Table 2**). Glatard et. al’s FSL-based analyses, namely the brain extraction (Brain Extraction Tool [FSL-BET]), tissue classification (FMRIB’s Automated Segmentation Tool [FSL-FAST]), image registration (FMRIB’s Linear Registration Tool [FSL-FLIRT]), and subcortical tissue segmentation (FMRIB’s Integrated Registration and Segmentation Tool [FSL-FIRST]) were replicated in each of these environments using 157 T1-weighted magnetic resonance images (MRI) from the International Consortium for Brain Mapping (ICBM)^108^. Each analysis was run twice within each environment to verify that there was no intra-environment variability. To evaluate the reproducibility of the analysis environment using locally installed vs. Neurodesk software, we compared the outputs for each installation type across computers (System A vs. System B). For intra- and inter-environment comparisons, we first compared file checksums. When two files produced different checksums, we quantified the pairwise differences across systems by computing Dice dissimilarity coefficients across images (**Figure 2a**). Note that there were never any intra-system differences in checksums (i.e., all analyses were deterministic, resulting in identical outcomes when run twice in the same computing environment). The code used to implement these analyses is available and re-executable through Neurodesk Play at: https://osf.io/e6pw3/.

**Figure 2.**
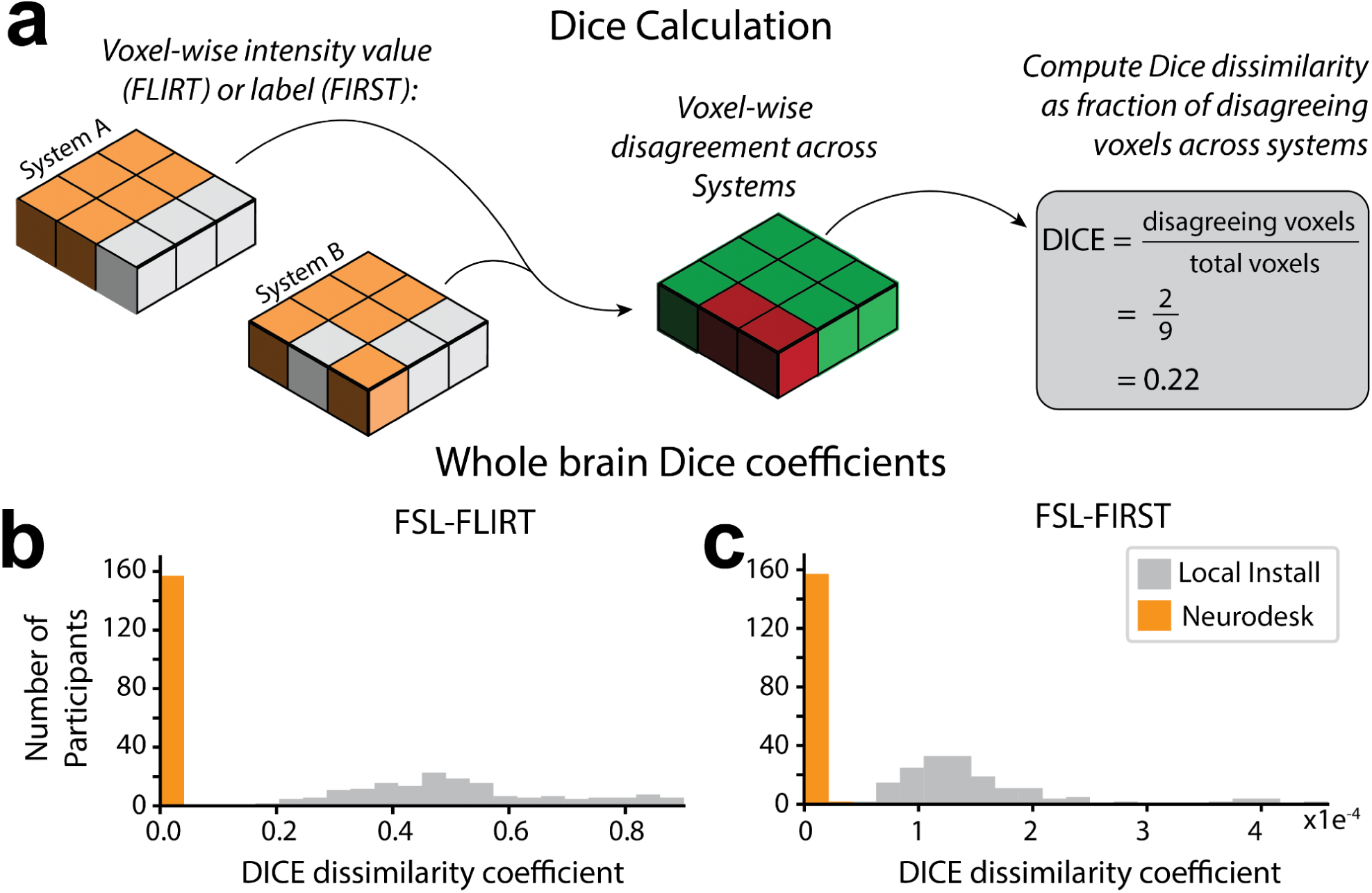
Discrepancies in image registration and tissue segmentation. (**a**) Calculation of the Dice dissimilarity coefficients; for each image, the voxel-wise disagreement in image intensity (FLIRT) or label (FIRST) calculated on System A vs System B was expressed as a proportion of the total number of voxels for each participant. (**b**) Histograms of Dice dissimilarity coefficients for image intensity calculated with FSL-FLIRT on Neurodesk vs. Local Install. To calculate these Dice coefficients, “disagreement” meant a voxel had a different intensity after image registration on System A vs. System B. Thus, the Dice coefficient of 0 for every participant whose images were registered using Neurodesk, means that the image intensity of each participant was matched across systems at every voxel. (**c**) Histograms of Dice dissimilarity coefficients for subcortical structure labels calculated using FSL-FIRST on Neurodesk vs. Local Install. To calculate these Dice coefficients, “disagreement” meant a voxel had different labels (e.g., amygdala, hippocampus, etc.) after image segmentation on System A vs. System B. Note that these Dice coefficients are much smaller than for image registration. This is expected because there are 73 times more “classes” for the image registration task, which uses image intensity (Range: 0 − 1903) as a class, than the classification task, which has labels for 15 structures. However, while both Neurodesk and the local system show strong agreement across systems overall, these distributions are completely non-overlapping, with Neurodesk showing much greater reliability across systems.

**Table 2.**
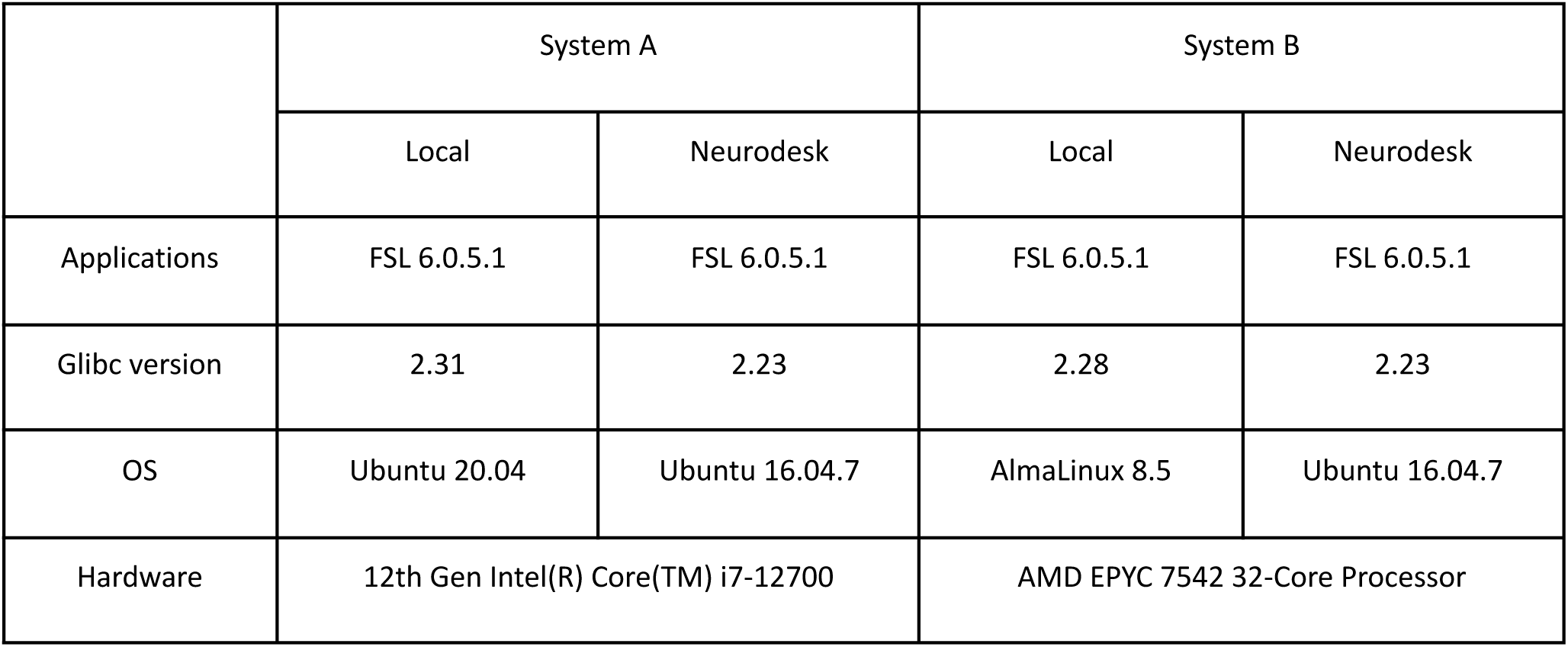
Computing environments used to run analyses.

|  | System A |  | System B |  |
| --- | --- | --- | --- | --- |
|  | Local | Neurodesk | Local | Neurodesk |
| Applications | FSL 6.0.5.1 | FSL 6.0.5.1 | FSL 6.0.5.1 | FSL 6.0.5.1 |
| Glibc version | 2.31 | 2.23 | 2.28 | 2.23 |
| OS | Ubuntu 20.04 | Ubuntu 16.04.7 | AlmaLinux 8.5 | Ubuntu 16.04.7 |
| Hardware | 12th Gen Intel(R) Core(TM) i7-12700 |  | AMD EPYC 7542 32-Core Processor |  |

#### Brain extraction and tissue classification

Skull stripping and brain tissue segmentation were done by running FSL BET and FAST on raw images. The pipeline was fully reproducible up to this stage because the file checksums were identical across all computing environments.

#### Image registration

FSL FLIRT was applied to register the images to the standard MNI-152 T1 1 mm template using 12 degrees of freedom. When run through Neurodesk, the outputs of this processing step had identical file checksums across computing systems for all images. However, file checksums for local installations of FSL did not match across systems. Dice dissimilarity coefficients for each image were computed to quantify the pairwise differences in image intensity across systems (**Figure 2a**). Voxel-wise agreement in image registration for Neurodesk was perfect (Dice dissimilarity coefficient; Range: 0.00, M = 0.00, SD = 0.00). However, there were many voxels with differing intensity across local installations (Dice dissimilarity coefficient; Range: 0.19 − 0.90, M = 0.51, SD =0.17, **Figure 2b**). These high Dice dissimilarity coefficients for the local installation indicate differences across many voxels, however, the magnitude of these differences in image intensity was subtle (inter-system intensity difference; M = 1.88, SD = 1.97; where *intensity ∈ Z: intensity ∈*[0, 1903], **Figure 3a, b**).

**Figure 3.**
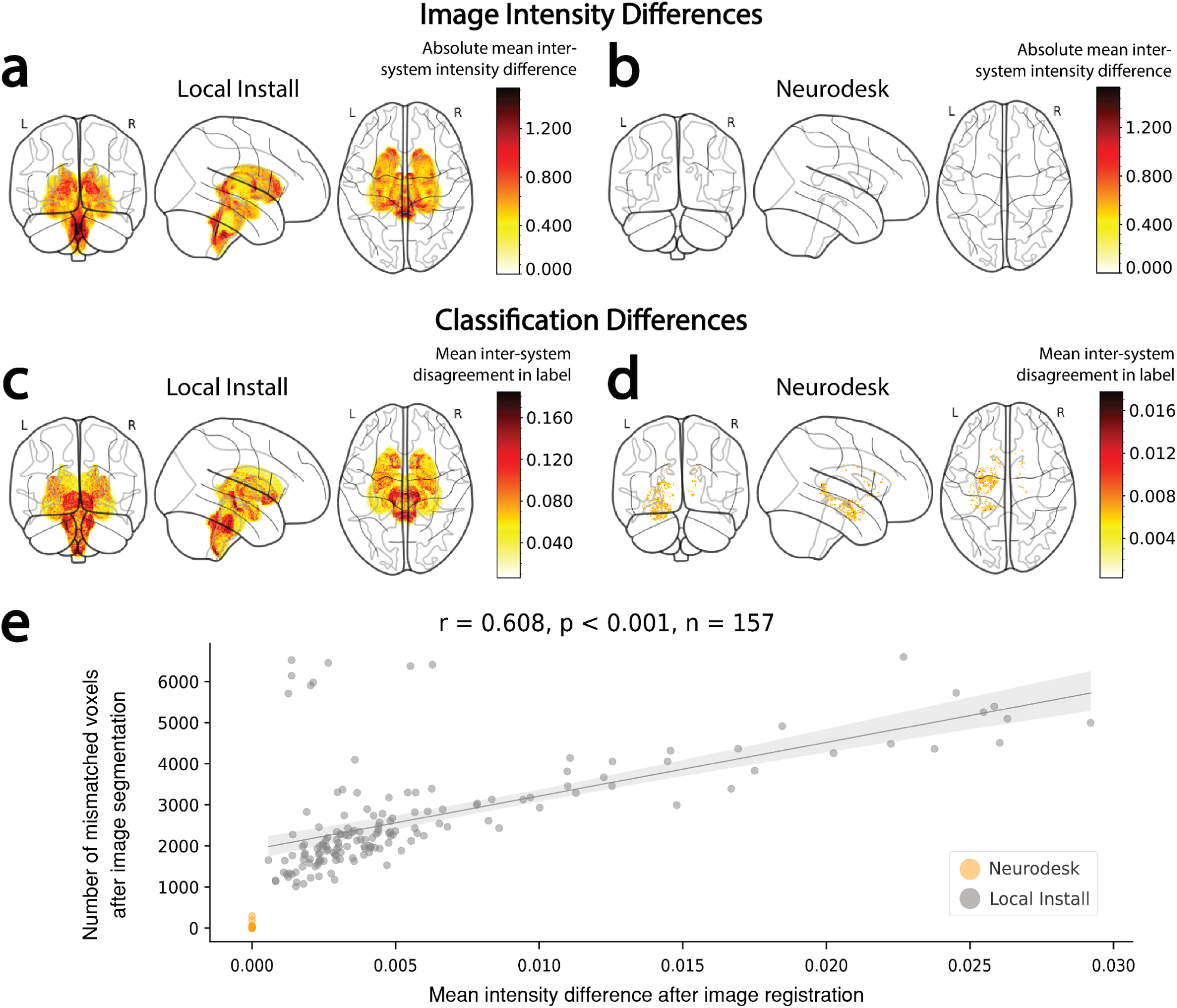
Inter-system differences in image intensity in subcortical structures and subsequent classification of these subcortical structures. (**a,b**) Absolute voxel-wise differences in image intensity within subcortical structures after image registration with FSL-FLIRT on each system (i.e.|Intensity_system A_ − Intensity_system B_|), averaged across participants. Projections are shown for image registration performed (**a**) using locally installed software, and (**b**) using Neurodesk (for which there were no intersystem differences). (**c,d**) Inter-system disagreement in subcortical structure labels after image segmentation with FSL-FIRST, averaged across participants. Projections are shown for image segmentation performed (**c**) using locally installed software and (**d**) using Neurodesk. (**e**) Scatter plot showing the mean inter-system image intensity differences across all voxels within the classified subcortical structures vs. the number of voxels subsequently classified with different labels across systems. For analyses performed with locally installed software, participants with larger differences in image intensity typically also had more prolific disagreement in labels between systems (Pearson’s r = 0.608, p < 0.001). This trend could not be assessed for Neurodesk, as there were no differences in image intensity across systems.

#### Subcortical tissue segmentation

Differences in image intensity across local installations were widespread yet subtle. In line with Glatard et. al’s approach, we next asked whether these differences impacted subcortical tissue segmentation (using FSL FIRST); the next step in the analysis pipeline. File checksums for the segmentation outputs matched for 0% of images when run using the local installation and for 93% of images when run with Neurodesk. Computation of the Dice dissimilarity coefficients for each type of installation revealed that while differences were small, they had non-overlapping ranges. Indeed, differences were much less prevalent for the Neurodesk installations (Dice dissimilarity coefficient; Range: 0.00 − 2.20×10^-5^, M = 3.43×10^-7^, SD < 0.01) compared with the local installations (Dice dissimilarity coefficient; Range: 5.80×10^-5^ − 4.59×10^-4^, M = 1.46×10^-4^, SD < 0.01, **Figure 2c**). On average, there were 426 times more voxel-wise disagreements across systems for the locally installed software than for Neurodesk. This difference can be visualized by comparing the 3D projections of the mean inter-system differences in classification across participants **(Figure 3c, d**). These projections illustrate that differences for locally installed software were widespread across all subcortical structures **(Figure 3c)**, while any subtle differences for Neurodesk were limited to a few voxels **(Figure 3d)**.

#### Understanding inter-system differences in image registration and tissue classification

Differences in tissue classification were at least partially attributable to differences in registered image intensity earlier in the pipeline. Indeed, there was a strong positive correlation between the magnitude of each participant’s inter-system differences in registered image intensity and inter-system classification mismatches (Pearson’s *r* = 0.608, *p* < .001, **Figure 3e)**. Thus, larger inter-system differences after the FSL FLIRT analysis were associated with larger inter-system differences after the subsequent FSL FIRST analysis.

We next replicated Glatard et al.’s findings by showing that the remaining variability in inter-system differences for tissue classification, as well as the differences for image registration, could be attributed to a combination of differences in floating point representation and differences in underlying dependencies across systems. Tracing the calls to dynamically linked system libraries revealed many differences for the local installations, but complete congruence between Neurodesk installations (**Figure 4**, see online methods). This begs the question - why were there still minor differences in the classification of subcortical structures for Neurodesk? The most likely explanation is that floating point calculations can produce different results on different processors due to different implementations of the floating point arithmetic instructions^109^. Reasons include whether 64 (SIMD, GPU) or 80 bit (x87 FPU) precision is used internally, reduced rounding for fused multiply-add, or if negative zero and positive zero are considered equal. Critically, these differences are minor, which is likely why the differences in classification across systems for Neurodesk were subtle.

**Figure 4.**
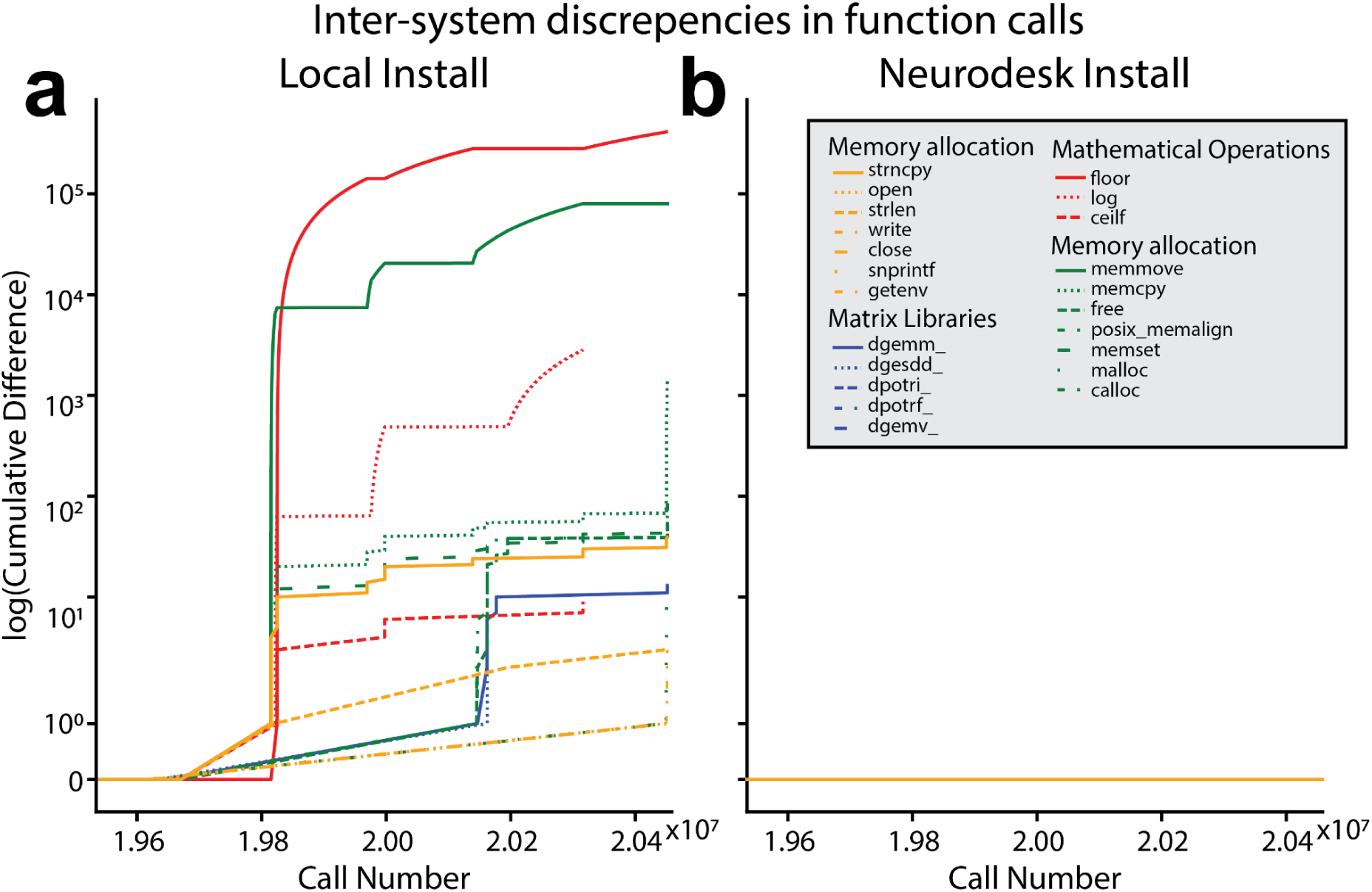
Cumulative difference in the numbers of system library calls between System A and System B for the analysis run using the (**a**) locally installed and (**b**) Neurodesk version of FSL FIRST. Note that calls to *floorf()* were excluded from the plot as they occurred earlier in time and the discrepancies for *floorf()* far outnumbered those for any other function from the locally installed tool.

Overall, these results demonstrate that differences in dependencies across computing environments can lead to slight differences in the outcomes of computational analyses. This can snowball across successive processing steps to cause potentially meaningful differences in results across computing environments, especially when investigating subtle effects. Minimizing differences at each stage of the analysis can enhance overall accuracy and reliability. Critically, Neurodesk eliminates this source of variability by facilitating access to containerized software. This allows researchers to reproduce the same result from different computing environments.

## Discussion

Neuroimaging data analysis pipelines are often challenged with limitations in *Accessibility*, *Flexibility*, *Portability* and *Reproducibility*. Neuroscientists may hold back from exploring new tools or spend excessive amounts of time installing software (and dependencies) in new computing environments, only to find that the same analysis pipeline produces different results. We developed Neurodesk to address these challenges by building an open-source and community-oriented platform for reproducible neuroimaging data analysis. Neurodesk allows scientists to flexibly create fully reproducible and accessible data analysis pipelines in various computing environments. By providing an accessibility layer for software containers, the Neurodesk platform allows for convenient portability across computing environments. Finally, by keeping the platform open-source and utilizing continuous integration and deployment, we have democratized the Neurodesk platform and set a path toward a sustainable ecosystem for neuroimaging data analysis.

The Neurodesk platform has the potential to transform neuroimaging data analysis, because it allows for truly reproducible data analysis and is highly accessible. Scientists strive to uphold scientific principles to the highest possible standard. However, looming deadlines and the pressure to publish often force individual researchers to find a balance between these ideals and the practical constraints imposed by resource limitations. Neurodesk can allow all researchers to adhere to the highest possible reproducibility standards with minimal changes to their typical development pipelines. Reducing unnecessary computational variability between execution systems makes it possible to share analyses between labs and collaborate on large datasets without potentially obscuring effects or introducing artificial differences between sites. Neurodesk enables researchers to not only access a comprehensive suite of neuroimaging data analysis software, but also contribute developments into the future for an ever-increasing suite of packages. Hence, researchers can flexibly take advantage of open datasets, reproduce reported analyses, switch between neuroimaging modalities across projects, and apply complementary analysis methods alongside their primary approach. By harnessing Neurodesk together with cloud computing technologies, published manuscripts can also include links to Jupyter Notebooks, therefore democratizing the reproducibility of key analyses. The ease with which Neurodesk allows analysis pipelines to be shared and reproduced across computing environments also has particular relevance for distributed research groups and collaborative, multi-site projects. Thus, the Neurodesk platform not only facilitates access to reproducible neuroimaging data analysis but also makes developing and sharing these workflows less burdensome.

Neurodesk is not the first platform to address the limited accessibility and reproducibility available for many neuroimaging data analysis tools. Indeed, software distribution mechanisms like NeuroDebian^2^ have made great progress in making neuroimaging software more accessible, while projects such as OpenNeuro^37^, Brainlife^38^, Flywheel^39^, XNAT^42^, Code-Ocean^110^, Boutiques^40,41^ and Qmenta^43^ have greatly improved the accessibility and reproducibility of neuroimaging analyses. However, all current existing solutions have lacked portability and flexibility. Many existing solutions require users to upload datasets to their platforms, and developing custom pipelines on these platforms requires substantial platform-specific knowledge. However, even users already accustomed to these specifics may still benefit from the Neurodesk project as Neurodesk’s containers are interoperable with other platforms.

Neurodesk has been developed as a research tool to facilitate the analysis of neuroimaging data. However, the platform may have a significant impact as an educational tool for workshops, summer schools, and ‘hackathons’^111^. The Neurodesk platform was first conceptualized during a ‘hackathon’ event, during which neuroscientists from around the globe gathered in hubs to collaborate on short-term projects, attend workshops, and develop critical research skills. One of the greatest hurdles for organizers and attendees of such events is the diversity in computing environments across researchers. When delivering a workshop or tutorial, facilitators often spend a large portion of the allocated time troubleshooting installations or issues specific to unique computing environments. Neurodesk addresses these issues by allowing access to identical computing environments with requisite tools pre-installed. This functionality allows groups of researchers to efficiently tackle complex problems by eliminating Sisyphean troubleshooting. The Galaxy platform, for example, has made a significant impact in this way by providing a containerized solution for bioinformatics and social science^112^. Aside from educational applications, Neurodesk can also aid research software developers wishing to make their tools more accessible. The effort to containerize and add one’s software to Neurodesk may be minimal compared to the burden of testing across multiple computing platforms and fielding support queries from end-users running software in diverse environments.

Neurodesk currently has limitations that warrant discussion. The first limitation is that the software containers in the Neurodesk platform currently do not support the ARM CPU architecture, which will become increasingly common as Mac users update their hardware. This stems from limitations in the underlying software applications, which currently need more support for this processor architecture. However, tool developers are rapidly adapting tools for this architecture, and we are convinced that this problem will be addressed for the most used applications in the future. Further limitations may arise as Neurodesk is applied across more diverse use-cases by the broader research community. A pertinent example relates to the use of proprietary and licensed software. This is an area of active development as the Neurodesk community investigates how to integrate such software without compromising the accessibility principle. A strength of Neurodesk is that the community-oriented, continuous integration model provides a powerful and flexible way to address such expanded use-cases without depending on a single development team. This relates to a potential limitation of any such platform - the project’s long-term sustainability. The Neurodesk platform was funded to be sustainable and supported by the community, but for this to be successful, the project needs constant maintenance. We, therefore, developed multiple pathways for sustainability, including the federated support of the underlying hosting infrastructure, flexibility in the continuous integration and deployment infrastructure, and a potential for a commercial model to offer tailored support for institutions and workshops.

The challenges to accessibility and reproducibility posed by neuroimaging data analysis software are not unique to neuroscience. While we have chosen to containerize software designed for neuroimaging datasets, the principles governing the design of the Neurodesk platform can be unrestricted to this field of research. This open-source platform could be used to deploy software specific to any other discipline, and it is our sincere hope that this platform is adapted to other disciplines struggling with similar issues. The Neurodesk platform has the potential to improve the way scientists analyze data and communicate results profoundly. For the first time, this platform allows any scientist, anywhere in the world, to conveniently access their data analysis tools and apply them in a fully reproducible manner from any computing environment. We are excited to see what new insights such technology can enable.

## Online methods

### Neurodesk’s open-access code and documentation

All stages of development, from the initial conception as a hackathon project, through to the most current iteration of Neurodesk, with up-to-date community-built Neurocontainer recipes, are documented publicly:

https://www.neurodesk.org/ - Platform website which includes ‘Getting Started’ tutorials for new users of various skill levels.

https://github.com/NeuroDesk - Public GitHub repository, where Issues can be logged, and contributions can be made by any community member with a GitHub account and the eagerness to create pull requests.

### Data Availability

The data that support the findings of this study are available from the International Consortium for Brain Mapping (ICBM) database (www.loni.usc.edu/ICBM). The ICBM project (Principal Investigator John Mazziotta, M.D., University of California, Los Angeles) is supported by the National Institute of Biomedical Imaging and BioEngineering. ICBM is the result of efforts of co-investigators from UCLA, Montreal Neurologic Institute, University of Texas at San Antonio, and the Institute of Medicine, Juelich/Heinrich Heine University - Germany. There are restrictions that apply to the availability of these data, which were used under approved permission for the current study, and so are not publicly available, but available from ICBM upon request.

### Code Availability

The code for this study is available on the GitHub repository at https://github.com/NeuroDesk with no restrictions on access. The code is licensed under the MIT License.

### Frequently Asked Questions

***How could researchers build an analysis pipeline and share this with other researchers using Neurodesk?***

We provide a Jupyter Notebook to showcase how different tools can be used in a fully reproducible and shareable analysis pipeline:

https://github.com/NeuroDesk/example-notebooks/blob/main/nipype_module_example.ipynb. In this example, we demonstrate the use of FSL and AFNI on a publicly available dataset. We used the open-source nipype workflow system to execute analyses on this data, enabling complex analyses to be built, shared, and executed identically in another Neurodesk installation.

***Will running my analyses on Neurodesk be slower than if they were run locally, especially if I’m on a slower internet connection?***

The internet bandwidth will only affect your analysis speed the first time you use a new tool. Neurodesk uses the CernVM File System (CVMFS), meaning that only the specific part of a currently used container will be downloaded over the internet. Once downloaded, these will be cached locally, meaning that software will operate at the same speed as it would when running locally (see **table S1**). Although there is a container initialization time that could impact performance in comparison to a non-containerized workflow, there is evidence that in some cases, containerized analysis pipelines may run even faster than locally installed software due to efficiency gains in accessing files^113^.

***Where are Neurodesk containers stored, and will the performance differ from country to country?***

Neurodesk containers are distributed globally via CVMFS and accessed from the fastest server according to your location. We aim to get mirror servers as close as possible to all users so that CVMFS can automatically use the fastest available mirror server.

***Are there any security concerns regarding using the Neurodesk platform in a web browser? For example, could there be any risks that compromise data processed on Neurodesk?***

The underlying container technology in Neurodesk ensures that applications are isolated with the least privileges to minimize the impact of malicious software. Interacting with the web from within a Neurodesktop poses a similar risk to any system with access to the internet, so all precautions would apply. Neurodesktop can be shut down, deleted and started fresh with minimal effort, which means recovery is significantly simpler than a native installation in a similar scenario. To ensure data security, it is essential for users who run Neurodesk on a cloud provider or in their local network to follow security best practices and secure the port Neurodesktop is running on via firewall rules. For an in-depth review of the potential security concerns of containerizing scientific data analysis software, see Kaur et al. (2021)^114^.

***Can I store processed data in Neurodesk?***

Neurodesktop allows host directories to be mounted for local data access, and these directories can then be accessed from the Neurocontainers. Data can also be accessed via access clients and the web inside a Neurodesktop instance running for example on a cloud provider. Upon installation of Neurodesk on a local PC or HPC, users have the option of mounting an existing local directory or utilizing the automatically created and locally stored directory, ∼/neurodesktop-storage. This directory is permanently stored on the local host and will remain even if Neurodesktop is deleted, ensuring that the data remains on the local host and does not leave their PC. It is important to note that the data remains on the user’s computer if Neurodesk is running locally, but Neurodesk can also run in a cloud environment where data is stored remotely and users need to ensure that their use case is in line with their ethics and data agreements.

***Can you provide more technical detail on how the Neurodesk desktop virtual environment has been built?***

Neurodesktop is a Docker container packaging a linux desktop environment that delivers neuroscience applications via CVMFS, distributed via singularity containers. It uses Apache Guacamole with underlying remote-desktop protocol (RDP) or virtual network computing (VNC) remote desktop protocols to deliver a desktop experience in the browser, including copy, paste and file transfer functionality.

***Why are there different types of containers (i.e. Docker, Singularity) in Neurodesktop? Are there any conflicts between Docker and Singularity?***

Docker and Singularity containers are both used in Neurodesktop for different, complementary purposes. Docker is used to containerize the Neurodesktop environment due its cross-platform support and ability to run singularity containers within. Singularity, which is used for the individual application containers (Neurocontainers), is preferred by most high-performance computing (HPC) platforms, where multi-user security and scheduling are of particular concern and can also be used indirectly via wrapper scripts and lmod; a system which manages environment configurations for different software packages.

***Are there any financial costs associated with keeping Neurodesk running, and if so, how will these be met for the foreseeable future?***

The long-term sustainability of Neurodesk has been planned according to three possible financial scenarios. *1) No further funding*: In this case, Neurodesk will be minimally maintained such that all the open-access containers will still be accessible. However, Neurodesk Play (the cloud-based no-install version of Neurodesktop) will no longer be accessible and the software distribution via CVMFS Neurodesk may run more slowly outside of Australia. *2) Marginal Funding.* Neurodesk will be maintained with its current functionality, but with less focus on the development of new features. *3) Sufficient funding*. The Neurodesk team is working on a not-for-profit business model in which additional financial costs involved in extending Neurodesk’s current functionality could be covered by charging a nominal fee to manage the resources required to deploy Neurodesk in combination with Jupyterhub in the cloud for organizations or for workshop and teaching purposes. Note that Neurodesk (Neurodesktop, Neurocommand, and the Neurocontainers) will always remain open-source and open-access under the MIT license, which enables commercial use. Any fee would be used to reduce the administrative load and technical challenges for workshop organizers and participants, such that workshop participants can access a fully maintained and cloud-based Neurodesktop environment.

***Neurodesk is open-source, such that anyone is able to contribute containerized software to the platform. Are there any protocols in place to verify that this software is working as expected before it is made available to the community?***

There is a feature to include a functional test within each tool’s container. This test can be run automatically after each container is built. However, such automated tests can only cover a subset of potential problems and we also rely on issues reported by users on GitHub and manual testing of new containers when releasing new versions.

***The software I need is not available in Neurodesk, and I don’t feel confident in my ability to contribute a container to the Neurodesk repository. Is there a way I can request that it be added?***

Users can submit a GitHub issue to request new tools by providing the following information: name and version of the tool, preferred Linux distribution, Linux commands for installing the tool, any GUI applications and commands to start them, test data to include, reference to paper, link to documentation, and commands for testing the tool.

***How do I get help if I encounter an issue with Neurodesk?***

There is an active discussion forum on GitHub with a Q&A section. If your question has not already been addressed there, please raise a new issue.

### Reproducibility in Neurodesk

To investigate our claims that the Neurodesk platform’s containerized tools lead to more reproducible results than locally installed software, we sought to conceptually replicate the results reported by Glatard et al. (2015) using Neurodesk vs locally installed software across different operating systems. The first steps in Glatard et al.’s analysis pipeline were brain extraction and tissue classification.

#### Brain extraction and tissue classification

FSL BET and FAST were run on raw MRI images to extract voxels containing brain tissue and classify tissue types, respectively. The file checksums for the outputs of these processing steps were identical across all computing environments, verifying that the implementation of the processing pipeline was reproducible across systems for both Neurodesk and local installation. After these steps, image registration and tissue classification were performed with FSL-FLIRT and FSL-FIRST, respectively. These analysis steps did lead to differences in results across systems, and are thus reported in the main text.

#### Understanding inter-system differences in image registration and tissue classification

Given that the image registration and tissue classification steps led to inter-system differences, we sought to understand the cause of these differences. FSL utilizes dynamic linking to shared system libraries such as libmath and LAPACK, which are loaded at runtime. Thus, while the same version of FSL was installed in all four computing environments, differences in image processing still emerge for analyses run on locally installed software. This is due to differences in dependencies across systems, a problem addressed by Neurodesk. To better understand how such differences might emerge, calls to these libraries were recorded for a representative image using ‘ltrace’. The libraries called during the FLIRT and FIRST analyses could be categorized into four main classes: mathematical operations, matrix operations, memory allocation, and system operations. Interestingly, Glatard et al., who used older software versions than we investigated here, found that image processing differences across systems resulted largely from differences in floating point representation in the mathematical functions *expf()*, *cosf()*, and *sinf()*. They also found inter-system differences in the control-flow of the programs, indicated by differences in the number of library calls to mathematical functions such as *floorf()*. Here, differences in floating point representation were less severe, as these were only present for the *sinf()* function. However, the number of calls made to several functions differed across the local FSL installations, indicating that the inter-system differences in the control flow of the processing pipeline remain an issue for reproducibility (**Table S1**). The *floorf()* function represented the most prevalent difference in library calls. There were over 13 000 additional calls to this function made on System B relative to System A for the FLIRT analysis, and approximately 5.5 million additional calls for the FIRST analysis. Overall, the FIRST analysis had greater discrepancy in calls overall. After accounting for the additional calls to *floorf()*, which occurred early in the FIRST analysis pipeline, mismatches in the sequence of system calls to several other functions remained (**Figure 4a**). However, all remaining mismatches across systems occurred in memory allocation functions. Importantly, there was no difference in floating point representation or the number of system calls to shared libraries across systems for the Neurodesk implementation of FSL (**Figure 4b**), while maintaining a similar runtime as local installation on the same hardware (**Table S1**).

**Table S1.**
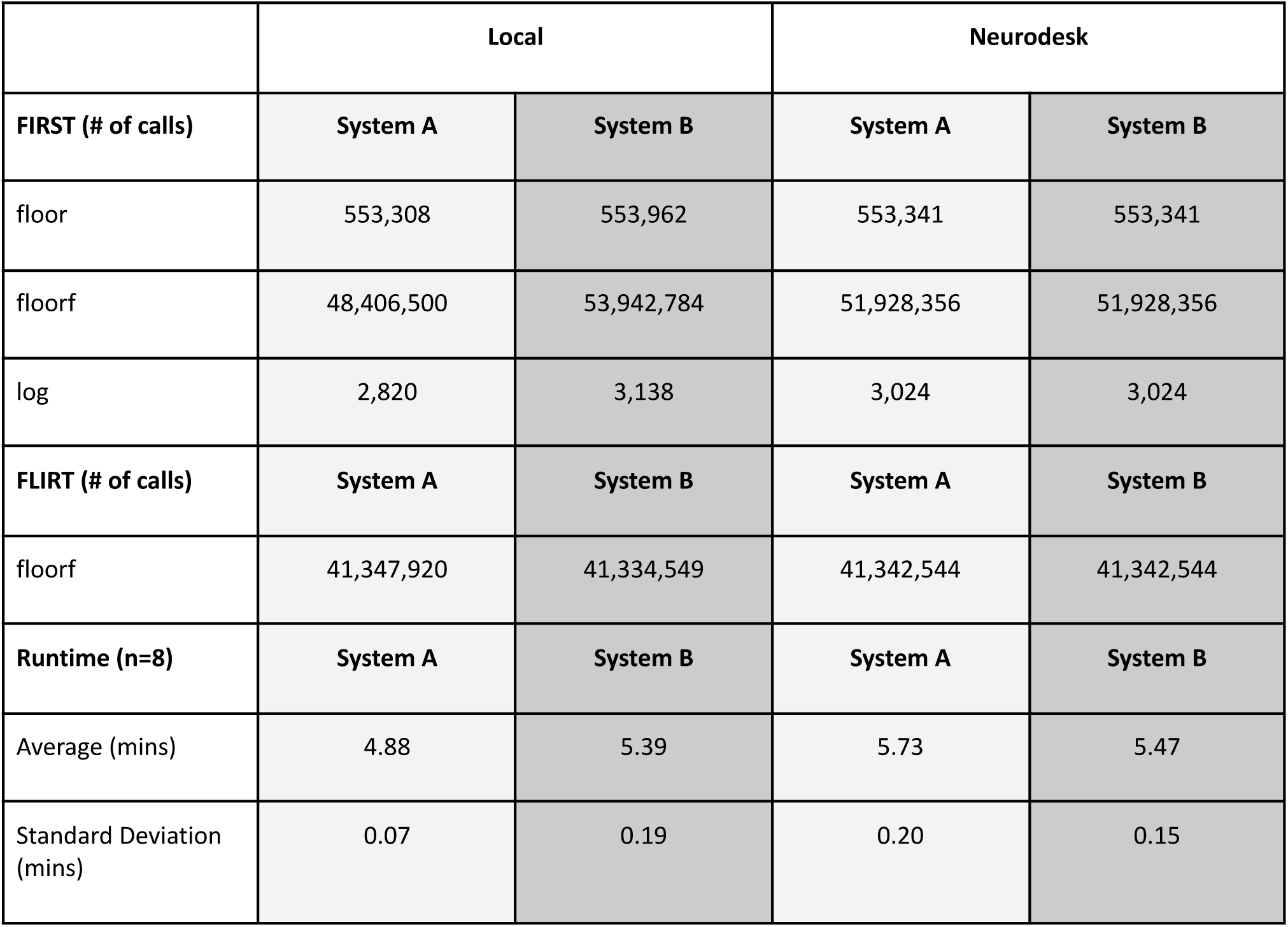
Differences in the execution of tissue segmentation (FIRST) and image registration (FLIRT) pipelines. Runtime refers to the CPU time spent on system and library calls within a pipeline.

#### Understanding the practical implications of inter-system differences

The local installations led to inter-system differences in tissue classification orders of magnitude larger than in Neurodesk. However, it is difficult to know how voxel-wise differences of this scale might actually affect test statistics i.e. could there actually be a different conclusion about the research question if the same analysis on the same data runs on a different computer? To address these questions, we performed a permutation test to examine the impact of inter-system differences in tissue classification (using FSL FIRST) on correlations between subcortical structure volumes and age.

On each system (A,B), for both Neurodesk and local installations, we computed the volume of each subcortical structure in the left hemisphere, right hemisphere, and the whole structure by participant. We performed permutation tests for each of these volumes (9999 permutations each). On each permutation, we performed a Pearson correlation of volume vs. participant age, and calculated the differences in the values of the correlation coefficients across the two systems. These permutation tests were repeated for three different sample sizes (n=10, 30, 50), such that each permutation for each sample size represented a different randomly selected group of participants. Critically, for each sample-wise permutation, the same sample was used for each of the two systems, such that the test-statistic difference always represented inter-system differences rather than inter-sample differences. Thus, the distribution of test statistic differences for each sample size represents 209979 permuted samples (7 subcortical structures (Putamen, Amygdala, Thalamus, Pallidum, Caudate Nucleus, Hippocampus, Accumbens.) x 3 methods (left hemisphere, right hemisphere, both) x 9999 subject-wise permutations).

The analysis showed that as sample size decreased, the inter-system coefficient differences for the local installations increased in magnitude (Local installation: N=50, Δr = -0.02 − 0.02 | N=30, Δr = -0.04 − 0.03 | N=10, Δr = -0.08 − 0.11; **Figure S1**). By contrast, the inter-system test statistic differences for Neurodesk were negligible and did not scale with sample size (Neurodesk: N=50, Δr = -1.74×10^-3^ − 2.59×10^-4^ | N=30, Δr = -3.75×10^-5^ − 1.89×10^-4^ | N=10, Δr = -1.52×10^-3^ − 0; **Figure S1**). Thus, the minor differences in image processing with locally installed software can meaningfully impact the reliability of test statistics, especially when statistical power is already low. It is therefore crucial to consider both sample variability and system variability when conducting these types of analyses.

**Figure S1.**
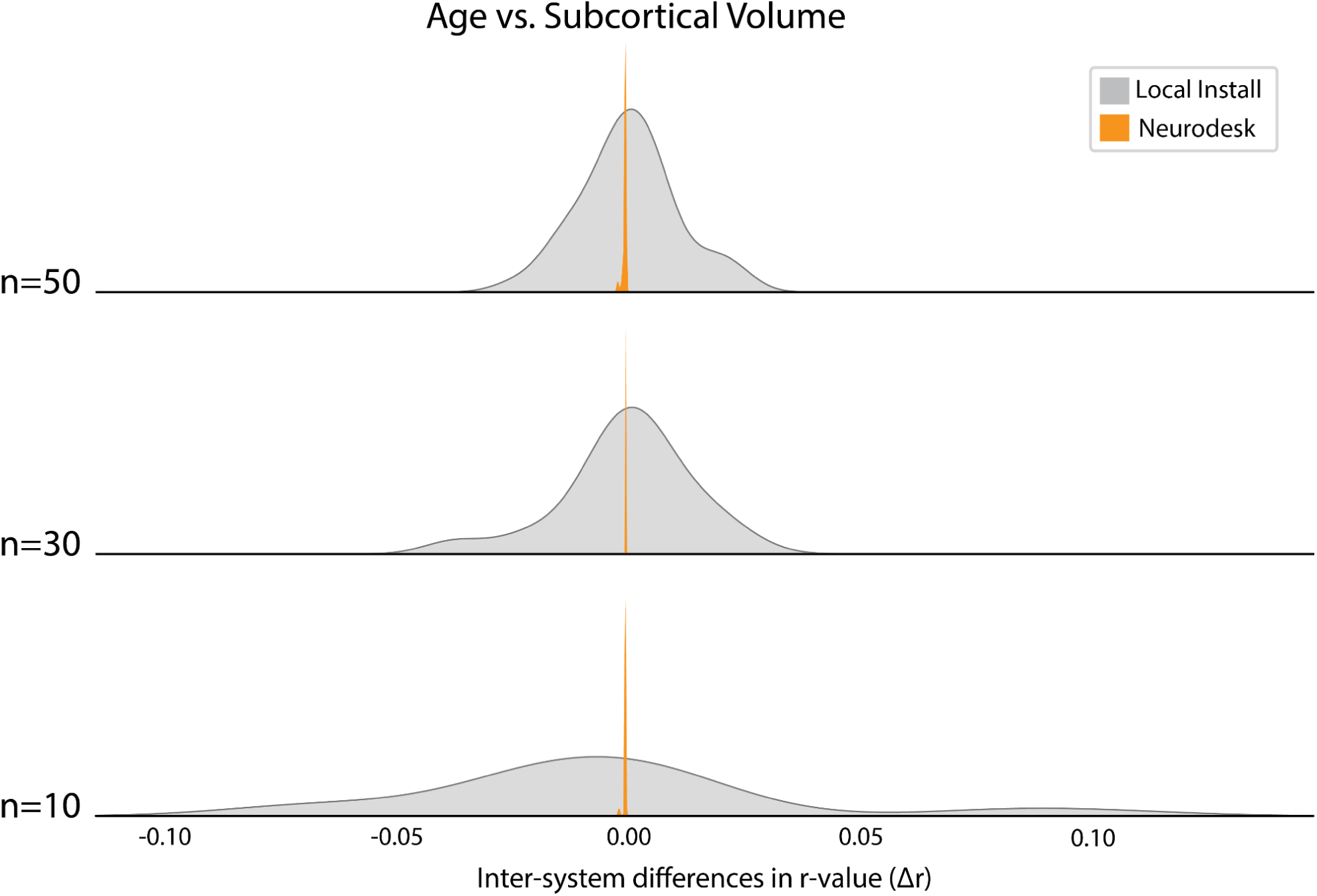
Permutation test results showing inter-system differences in r-values for the correlation between age and volume of subcortical structures, organized by sample size (n = 10, 30, 50).

## Acknowledgements

The Australian Research Data Commons (ARDC) invested in Neurodesk’s development through the Australian Electrophysiology Data Analytics Platform (AEDAPT) project. Thank you to Oracle for Research for providing Oracle Cloud credits and related cloud resources to support this project. The University of Queensland funded the project via the Knowledge Exchange & Translation (Kx&T) Fund and the UQ AI Collaboratory. The authors acknowledge the facilities and scientific and technical assistance of the National Imaging Facility, a National Collaborative Research Infrastructure Strategy (NCRIS) capability. NIH P41EB019936 partially supported JRK and SSG. Data collection and sharing for this project was provided by the International Consortium for Brain Mapping (ICBM; Principal Investigator: John Mazziotta, MD, PhD). ICBM funding was provided by the National Institute of Biomedical Imaging and BioEngineering. ICBM data are disseminated by the Laboratory of Neuro Imaging at the University of Southern California.

## Author contributions (CRediT statement^1^)

***Conceptualization*:** Steffen Bollmann, Aswin Narayanan, Oren Civier, Tom Johnstone, David White, Angela Renton, Thomas Shaw, Ryan Sullivan, Thomas Close, Anthony Hannan, Gary Egan, Marta Garrido, Aina Puce, Franco Pestilli, Martin Grignard, Levin Kuhlmann, Gershon Spitz, David Abbott, Megan Campbell, Nigel Rogasch, Jakub Kaczmarzyk, Satrajit Ghosh. ***Software*:** Steffen Bollmann, Aswin Narayanan, Thomas Shaw, Oren Civier, Tom Johnstone, David White, Akshaiy Narayanan, Thuy Dao, Ashley Stewart, Martin Grignard, Lars Kasper, Judy D Zhu, Korbinian Eckstein, Stefanie Evas, Xincheng Ye, Fernanda Ribeiro, Jeryn Chang, Kexin Lou, Jo Morris, Renzo Huber, Yorguin-Jose Mantilla-Ramos, Andy Botting, Chris Rorden. ***Validation*:** Steffen Bollmann, Aswin Narayanan, Toluwani J. Amos, Angela Renton, Thomas Shaw, Oren Civier, David White, Kelly Garner, Thuy Dao, Ashley Stewart, Lars Kasper, Judy D Zhu, Korbinian Eckstein, Guillaume Flandin, Martin Grignard, Stefanie Evas, Xincheng Ye, Mark Schira, Fernanda Ribeiro, Jeryn Chang, Jakub Kaczmarzyk, Kexin Lou, Renzo Huber, Ryan Sullivan, Thomas Close, Matthew Hughes, Levin Kuhlmann, Gershon Spitz, David Abbott, Megan Campbell, Nigel Rogasch, Marta Garrido, Aina Puce. ***Formal analysis*:** Thuy Dao. ***Conceptualization of Formal analysi*s:** Steffen Bollmann, Thuy Dao, Angela Renton, Fernanda Ribeiro, Thomas Shaw. ***Writing - Original Draft*:** Angela Renton. ***Writing - Review & Editing*:** All Authors. ***Writing - Initial Outline***: Angela Renton, Oren Civier, Paris Lyons, Steffen Bollmann. ***Visualization*:** Angela Renton. ***Supervision*:** Steffen Bollmann, Tom Johnstone, Angela Renton. ***Project administration*:** Steffen Bollmann, Aswin Narayanan, Paris Lyons, Tom Johnstone, Oren Civier, Benjamin Slade. ***Funding acquisition*:** Steffen Bollmann, Aswin Narayanan, Oren Civier, Tom Johnstone, David White, Ryan P. Sullivan.

