## Supplementary material for "Neurodesk: An accessible, flexible, and portable data analysis environment for reproducible neuroimaging": Online Methods

**Table S1.** Differences in the execution of tissue segmentation (FIRST) and image registration (FLIRT) pipelines. Runtime refers to the CPU time spent on system and library calls within a pipeline.

| | Local | | Neurodesk | |
|---|---|---|---|---|
| **FIRST (# of calls)** | **System A** | **System B** | **System A** | **System B** |
| floor | 553,308 | 553,962 | 553,341 | 553,341 |
| floorf | 48,406,500 | 53,942,784 | 51,928,356 | 51,928,356 |
| log | 2,820 | 3,138 | 3,024 | 3,024 |
| **FLIRT (# of calls)** | **System A** | **System B** | **System A** | **System B** |
| floorf | 41,347,920 | 41,334,549 | 41,342,544 | 41,342,544 |
| **Runtime (n=8)** | **System A** | **System B** | **System A** | **System B** |
| Average (mins) | 4.88 | 5.39 | 5.73 | 5.47 |
| Standard Deviation (mins) | 0.07 | 0.19 | 0.20 | 0.15 |

*Understanding the practical implications of inter-system differences.* The local installations led to inter-system differences in tissue classification orders of magnitude larger than in Neurodesk. However, it is difficult to know how voxel-wise differences of this scale might actually affect test statistics i.e. could there actually be a different conclusion about the research question if the same analysis on the same data runs on a different computer? To address these questions, we performed a permutation test to examine the impact of inter-system differences in tissue classification (using FSL FIRST) on correlations between subcortical structure volumes and age.

On each system (A,B), for both Neurodesk and local installations, we computed the volume of each subcortical structure in the left hemisphere, right hemisphere, and the whole structure by participant. We performed permutation tests for each of these volumes (9999 permutations each). On each permutation, we performed a Pearson correlation of volume vs. participant age, and calculated the differences in the values of the correlation coefficients across the two systems. These permutation tests were repeated for three different sample sizes (n=10, 30, 50), such that each permutation for each sample size represented a different randomly selected group of participants. Critically, for each sample-wise permutation, the same sample was used for each of the two systems, such that the test-statistic difference always represented inter-system differences rather than inter-sample differences. Thus, the distribution of test statistic differences for each sample size represents 209979 permuted samples (7 subcortical structures (Putamen, Amygdala, Thalamus, Pallidum, Caudate Nucleus, Hippocampus, Accumbens.) x 3 methods (left hemisphere, right hemisphere, both) x 9999 subject-wise permutations).

The analysis showed that as sample size decreased, the inter-system coefficient differences for the local installations increased in magnitude (Local installation: N=50, $\Delta r$ = -0.02 – 0.02 | N=30, $\Delta r$ = -0.04 – 0.03 | N=10, $\Delta r$ = -0.08 – 0.11; **Figure S1**). By contrast, the inter-system test statistic differences for Neurodesk were negligible and did not scale with sample size (Neurodesk: N=50, $\Delta r = -1.74 \times 10^{-3}$ – $2.59 \times 10^{-4}$ | N=30, $\Delta r = -3.75 \times 10^{-5}$ – $1.89 \times 10^{-4}$ | N=10, $\Delta r = -1.52 \times 10^{-3}$ – 0; **Figure S1**). Thus, the minor differences in image processing with locally installed software can meaningfully impact the reliability of test statistics, especially when statistical power is already low. It is therefore crucial to consider both sample variability and system variability when conducting these types of analyses.

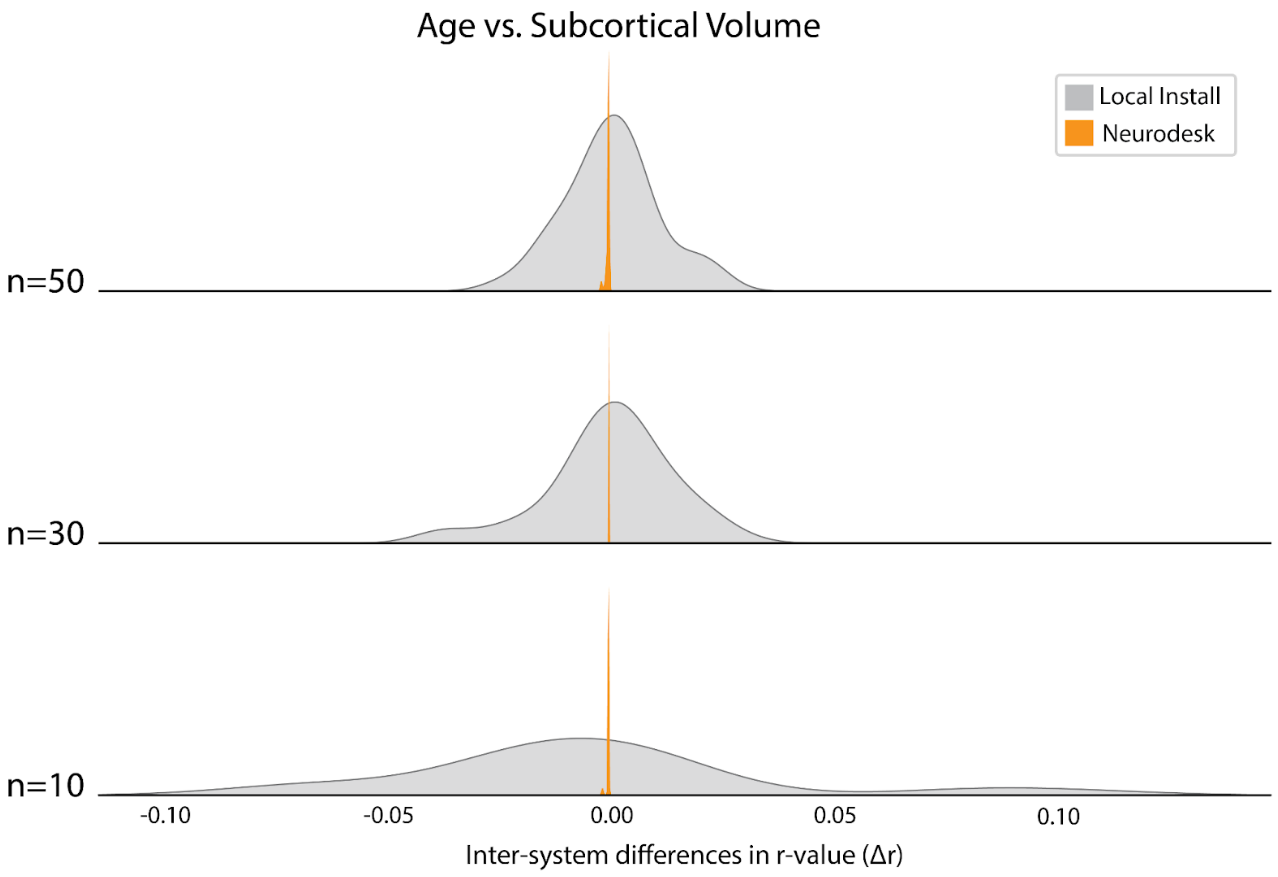


**Figure S1.** Permutation test results showing inter-system differences in r-values for the correlation between age and volume of subcortical structures, organized by sample size (n = 10, 30, 50).
